# Host adaptation and microbial competition drive *Ralstonia solanacearum* phylotype I evolution in South Korea

**DOI:** 10.1101/2020.07.10.196865

**Authors:** Maxim Prokchorchik, Ankita Pandey, Hayoung Moon, Wanhui Kim, Hyelim Jeon, Stephen Poole, Cécile Segonzac, Kee Hoon Sohn, Honour C. McCann

## Abstract

Bacterial wilt caused by the *Ralstonia solanacearum* species complex (RSSC) threatens the the cultivation of important crops worldwide. The exceptional diversity of type III secreted effector (T3E) families, high rates of recombination and broad host range of the RSSC hinder sustainable disease management strategies. We sequenced 30 phylotype I RSSC strains isolated from pepper (*Capsicum annuum*) and tomato (*Solanum lycopersicum*) in South Korea. These isolates span the diversity of phylotype I, have extensive effector repertoires and are subject to frequent recombination. Recombination hotspots among South Korean phylotype I isolates include multiple predicted contact-dependent inhibition loci, suggesting microbial competition plays a significant role in *Ralstonia* evolution. Rapid diversification of secreted effectors present challenges for the development of disease resistant plant varieties. We identified potential targets for disease resistance breeding by testing for allele-specific host recognition of T3Es present among South Korean phyloype I isolates. The integration of pathogen population genomics and molecular plant pathology contributes to the development of location-specific disease control and development of plant cultivars with durable resistance to relevant threats.

**Repositories:** All genome sequences obtained in this study are deposited to NCBI GeneBank under BioProject number PRJNA593908

## Introduction

*Ralstonia solanacearum* is a soil-borne pathogen that causes bacterial wilt disease of over 200 species in 50 different plant families, presenting a major threat to global crop production. This β-proteobacterium colonizes vascular tissues by invading root systems via wounds. Rapid proliferation and biofilm formation block water flow, causing wilt and death [1, 2]. *R. solanacearum* causes severe outbreaks in crops of critical economic and nutritional importance, including banana *(Musa* spp.), potato (*Solanum tuberosum*), tomato (*S. lycopersicum*) and pepper (*Capsicum annuum*) [1]. The broad host range, extensive distribution and genetic heterogeneity of *R. solanacearum* present major obstacles for the design of sustainable disease management strategies.

*R. solanacearum* was initially described as a single species within *Burkholderia* prior to reassignment to *Ralstonia* [3, 4]. Low (< 70%) levels of DNA hybridization between strains indicated *R. solanacearum* is more appropriately described as a species complex (RSSC) [7]. Four RSSC phylotypes were delineated based on 16S–23S rRNA gene intergenic spacer (ITS) regions and the *hrpB* and *egl* genes, coupled with comparative genomic hybridization analysis. Phylotype I is mainly present in Asia, phylotype II mainly in the Americas, phylotype III in Africa and phylotype IV in Japan, Indonesia, Australia and the Philippines [1]. Multi-locus sequence analysis and whole-genome comparisons indicate phylotype I and III are more closely related to each other, while phylotype IV is most divergent. Although we refer to phylotype designations in this paper, taxonomic revisions have been proposed to reclassify the RSSC into three separate species: *Ralstonia pseudosolanacearum* (phylotypes I and III), *R. solanacearum* (phylotype II) and *R. syzygii* (phylotype IV) [5–7].

*R. solanacearum* phylotypes have ∼ 5 Mb bipartite genomes comprised of two replicons 3.5-3.7 Mb and ∼ 1.6-2.1 Mb in size. The smaller replicon is conventionally referred to as a megaplasmid, though both replicons encode essential and pathogenesis-related genes. Multipartite genomes are frequently observed among genera harbouring soil and marine bacteria interacting with eukaryotic hosts [8]. The mosaic structure of RSSC genomes attests to the importance of recombination in enhancing the genetic variation evident in this species complex [7, 11]. Natural competence is considered a ubiquitous trait among RSSC phylotypes and has been demonstrated both *in vitro* and *in planta* [9, 10]. Coupat et al. (2008) found 80% of 55 RSSC strains tested developed competence *in vitro* and exhibited single or multiple homologous recombination events up to 90 kb in length. High rates of recombination enhance the potential for rapid adaptation and emergence of novel pathogen variants [11, 12].

Virulence mechanisms in the RSSC include biofilm production, secretion of cell wall degrading enzymes and type III secreted virulence effector (T3E) proteins. Coordinated synthesis of T3Es is achieved by the activity of HrpG/HrpB transcriptional regulators on the hypersensitive response and pathogenicity (*hrp*) box cis-element in the promoter of T3E genes. RSSC T3Es target various compartments and functions to suppress host resistance and promote bacterial proliferation [13]. The RSSC has a vastly expanded type III effector repertoire compared to other well-studied plant pathogenic bacteria, with over one hundred T3E families identified thus far [14]. In contrast, 70 and 53 orthologous effector families are reported in *Pseudomonas syringae* and *Xanthomonas* spp. respectively [15, 16].

Plants have evolved intracellular receptors that monitor for the presence of pathogen effectors and activate a robust defense response often culminating in a form of programmed cell death (referred to as a hypersensitive response) around the site of infection and recognition [17]. Effector-dependent host specific interactions have been identified in *R. solanacearum*. A comprehensive survey of *R. solanacearum* phylotype II BS048 effector recognition identified RipAA, RipE1 and RipH2 as cell death inducers in *Nicotiana* spp., tomato and lettuce respectively [18]. The phylotype I GMI1000 RipE1 allele also triggers hypersensitive response in *Nicotiana benthamiana* [19, 20]. RipA family AWR motif-containing effectors are strong inducers of cell death in *Nicotiana* spp. [21]. The loss of RipAA was recently found to be correlated with the emergence of phylotype IIB strains pathogenic on cucurbits; related strains retaining RipAA are pathogenic on banana but not cucurbits [22]. We sequenced 30 RSSC phylotype I isolates from tomato and capsicum to identify the population structure and virulence factor repertoire of pathogens limiting production of these key crops in South Korea. Location and host of isolation do not have a strong impact on the population structure of South Korean phylotype I isolates; diverse phylotype I isolates capable of infecting solanaceous crops are broadly distributed across South Korea. Loci linked with competition are frequent targets of recombination, suggesting microbial interactions play an important role in *Ralstonia* evolution. We examined South Korean phylotype I virulence factor evolution in the context of the global pool of RSSC T3Es and found that most strains carry large effector repertoires with evidence of frequent horizontal transfer. We identified T3Es that elicit stable host recognition and defense responses regardless of which allelic variant is used, indicating they would be suitable targets for disease resistance breeding efforts. We also identified allelic variation in two T3Es resulting in the loss of host recognition. Rapid diversification of secreted effectors present challenges for the development of resistant plant varieties. The integration of pathogen population genomics with molecular plant pathology contributes to location-specific pathogen control strategies, as well as the development of cultivars with durable resistance to relevant threats.

## Methods

### Genome assembly and annotation

*Ralstonia solanacearum* (*Rso*) were isolated from field-grown pepper and greenhouse tomato plants between 1999 and 2008 (Table 1). Tomato isolates were collected from the north in 2008, while pepper isolates were collected from central and southern regions of the South Korean peninsula. One isolate (Pe_4) was collected from Jeju island, southeast of the peninsula. Genomic DNA was extracted using the Promega Wizard Genomic DNA Purification kit and 150 bp paired-end sequencing was run on the Illumina NextSeq platform using Nextera and the library preparation protocol described by Baym et. al (2015) [23]. Reads were filtered to remove adapter sequences (parameters: ktrim=r k=23 mink=11 hdist=1 minlen=50 ftm=5 tpe tbo) and common sequence contaminants according to NCBI Univec database (parameters: k=31 hdist=1), and quality-trimmed using the bbduk program available in the BBmap package (paramters: qtrim=r trimq=10 minlen=50 maq=10 tpe). Filtered and quality-trimmed reads were used for genome assembly with SPAdes v3.10.0, using --careful assembly with kmer size distribution –k 21, 33, 55 and 77 [24]. The coverage cutoff was set to auto. Coverage was assessed by mapping reads to the assembly using Bowtie2, removing duplicates [25]. Assembly improvement was performed using Pilon v1.14 and assembly statistics obtained using assembly-stats (https://github.com/sanger-pathogens/assembly-stats) [26]. Genomes were annotated with Prokka v.1.12 beta using a custom annotation database created using all 71 *Ralstonia solanacearum* annotated assemblies available on Genbank Nov. 11, 2017 [27]. 68 publicly available *R. solanacearum* genomes were reannotated with Prokka v1.13.3 as well for inclusion in downstream analyses (Table S1).

**Table 1.**
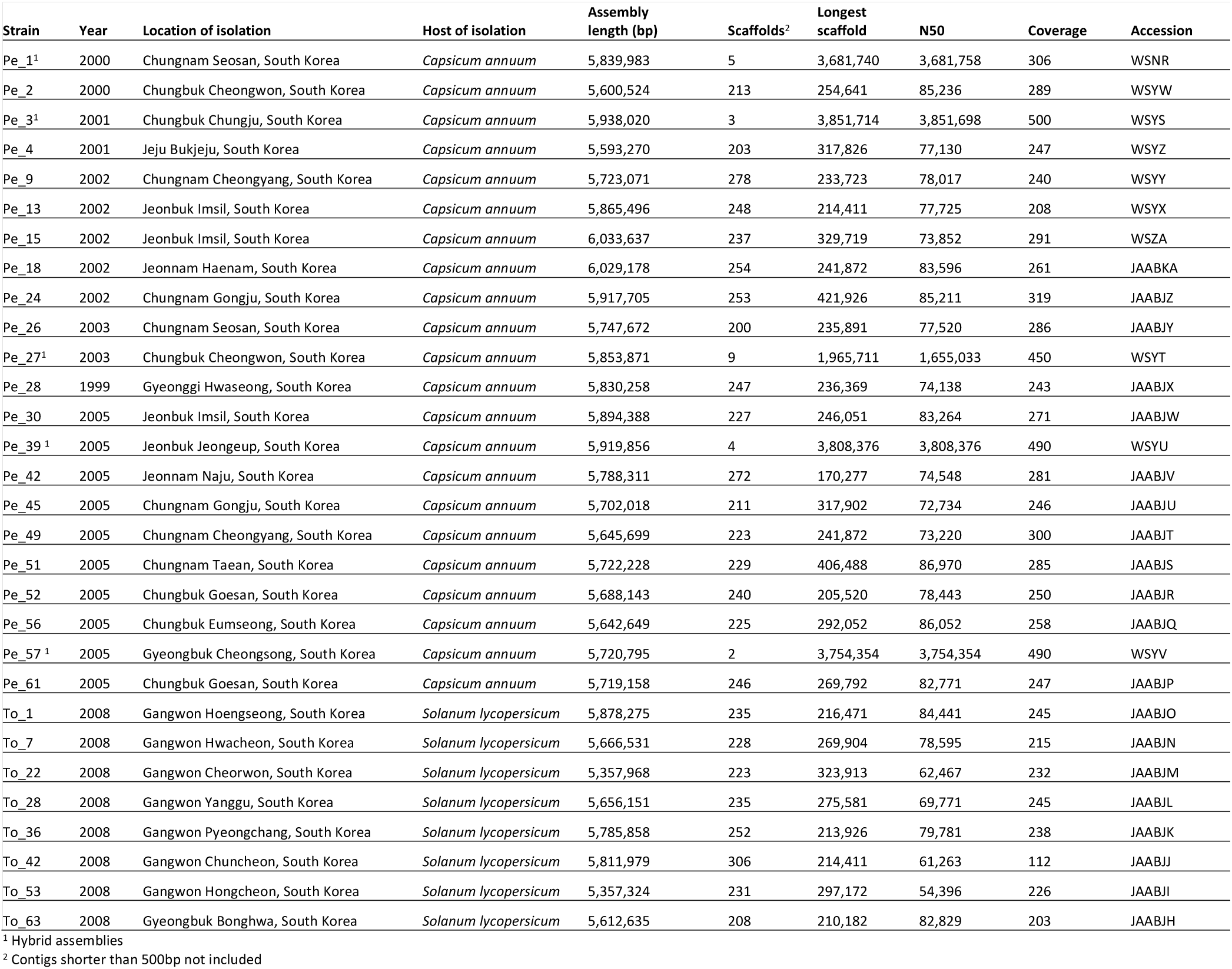
*R. solanacearum* phylotype I isolates and assembly statistics

Nanopore sequencing was performed for five isolates: Pe_1, Pe_3, Pe_27, Pe_39 and Pe_57. DNA extraction was performed using Promega Wizard, and barcoding and library preparation were performed using Native Barcoding Expansion 1-12 (EXP-NBD104) kit, in conjugation with the Ligation Sequencing Kit (SQK-LSK109) according to manufacturer recommendations. After sequencing run of 48 h, basecalling was performed using Guppy followed by hybrid assembly with Unicycler using Illumina reads for each respective strain [28]. Genomes were annotated with Prokka v1.13.3 using the custom *R. solanacearum* annotation database.

### Core genome phylogeny of Korean phylotype I isolates

The core genome phylogeny of South Korean phylotype I isolates was obtained by mapping Illumina reads to the completed nanopore assembly of *R. solanacearum* Pe_57. Regions of the reference genome corresponding to mobile genetic elements were removed after identification using the PHASTER phage and ISFinder databases; no integrative and conjugative elements were identified with ICEberg [29–31].

Readmapping of filtered, quality-trimmed reads was performed using BWA mem 0.7.17 - r1188 [32]. Samtools v1.3.1 was used to sort and index reads, remove duplicate reads and generate pileup data [31]. Variants were called with BCFtools v1.3.1 (bcftools mpileup -- consensus-caller --ploidy 1 --min-MQ 60) and a set of filtered SNPs obtained using BCFTools v0.1.13 to retain single nucleotide polymorphisms (SNPs) with a minimum read depth of 10 in both directions, non-reference allele frequency of 0.95, minimum quality of 30 and minimum distance from a gap of 10bp [33]. BEDtools v2.26.0 was used to determine positions where the depth of coverage fell below 10 reads [34]. A final consensus relative to the reference was obtained with BCFtools consensus v1.3.1, masking low coverage positions identified by BEDtools, retaining both invariant sites and SNPs passing coverage, non-reference allele frequency and quality thresholds. An alignment was generated using the consensus sequences of each isolate. Alignment positions with one or more gaps or overlapping mobile genetic elements in the reference genome were removed. Initial tree building was performed with RAxML v7.2.8 using parameters: -f a -p $RANDOM –x $RANDOM -# 100 -m GTRGAMMA -T 8 [35].

ClonalFrameML v1.178 was employed to identify polymorphisms introduced via homologous recombination using the gap-free core genome alignment and initial maximum likelihood phylogeny as input [36]. Alignment regions identified as likely to have been introduced via recombination were removed from the alignment and tree building was repeated using the non-recombinant alignment in RAxML v7.2.8 (parameters as above). Trees were visualized using FigTree v1.4.3, displaying only nodes with bootstrap support scores of 70 and above (https://github.com/rambaut/figtree/). This approach was employed to obtain a core genome phylogeny and alignment using the primary and secondary replicons of *R. solanacaerum* Pe_57.

Recent and ancestral recombination events among South Korean strains phylotype I strains were visualized using maskrc-svg.py with the core genome alignment along with the recombinant regions predicted by ClonalFrameML (https://github.com/kwongj/maskrc-svg). Recombination events are referred to as ancestral when predicted to occur at internal (ancestral) nodes, and extant or recent events when impacting terminal nodes (individual strains). The number of recent recombinant events was calculated with pyGenomeTracks by counting the number of extant (recent) recombination events overlapping with annotated coding sequences in the reference genome GMI1000 chromosome and megaplasmid, excluding recent events in GMI1000 and Pe_57 (https://github.com/deeptools/pyGenomeTracks).

### *R. solanacearum* species complex phylogeny

The *R. solanacearum* species complex tree was generated using an alignment of concatenated single-copy core genes. All Korean isolates and 68 publicly available *R. solanacearum* genomes were annotated with Prokka (as above). OrthoFinder was implemented to identify single copy orthologs [37]. Single copy ortholog nucleotide sequences were employed for codon-aware alignment with PRANK [38]. Core gene alignments were concatenated and stripped of gap and invariant positions. Maximum likelihood phylogeny reconstruction was then performed with RAxML v8.2.10 (parameters: -f a -p $RANDOM -x $RANDOM -# 100 -m GTRCATX -T 16) [36]. Trees were visualized using FigTree v1.4.3, displaying only nodes with bootstrap support scores of 70 and above (https://github.com/rambaut/figtree/). Estimation of the population structure was performed with RhierBAPS using the core SNP alignment with two levels of clustering and setting the number of initial clusters to 20 [39].

### Type III effector identification and comparative analyses

Effector predictions were performed on the South Korean phylotype I isolates and 68 publicly available genomes. First, sequences from the manually curated *R. solanacearum* effector (RalstoT3E) database were downloaded from (http://iant.toulouse.inra.fr/bacteria/annotation/site/prj/T3Ev3) and used as queries in tBLASTn searches of the 30 Korean *R. solanacearum* genomes, retaining all hits with a minimum coverage of 30% and pairwise identity of 50%. Overlapping hits on the subject sequence were merged to generate the longest hit using BEDtools. The merged hit was then scanned to identify the predicted open reading frame (ORF), searching for start and stop codons and the presence of any premature stop codons. The presence of putative *hrp* boxes up to 500 bp upstream of the effector candidate ORF start codon was retained. Merged hits containing no start codons and/or premature stop codons were adjusted to reconstitute a functional ORF if possible. Start codon adjustment was performed by searching between the merged hit and putative *hrp* box (or up to 500 bp upstream of the merged hit for a start codon). Effector candidates were classified as pseudogenes in the absence of a start codon or in the presence of a stop codon resulting in a truncated ORF 30% shorter than the shortest query sequence generating the initial hit.

Individual effector phylogenies were generated using PRANK codon-aware alignments followed by tree building using RAxML 8.2.12. Gene tree topologies were compared with the species complex core gene phylogeny using the normalized Robinson-Foulds metric implemented in ETE3, collapsing identical sequences into the same leaf and disregarding comparison between nodes with bootstrap scores lower than 70 [40].

### Bacterial strains and plant growth conditions

*Escherichia coli* DH5α and *Agrobacterium tumefaciens* AGL1 were grown in Luria-Bertani (LB) medium supplied with appropriate antibiotics at 37°C and 28°C, respectively. *Nicotiana benthamiana* and *N. tabacum* W38 plants were grown at 25°C in long-day conditions (14 h light/10 h dark).

### Plasmid construction for transient expression of effectors

*R. solanacearum* genomic DNA was isolated using the Promega Wizard bacterial genomic DNA extraction kit following the manufacturer’s protocol. Allelic variants of RipA1 and RipAA were cloned from South Korean phylotype I isolates as well as GMI1000 and BS048 using Golden Gate [41]. Alleles were PCR amplified using Takara DNA polymerase (Thermo) and cloned into pICH40121 as ∼1 kb modules with *Sma*I/T4 ligase (New England Biolabs). The modules in pICH40121 were assembled into binary vector pICH86988 under control of the constitutive 35S cauliflower mosaic virus promoter and in C-terminal fusion with 6xHA or 3xFLAG epitope. The binary constructs were mobilized into *Agrobacterium tumefaciens* strain AGL1 using electroporation for transient expression experiments in *Nicotiana* spp.

### Hypersensitive response and ion leakage assays

*A. tumefaciens* AGL1 carrying effector constructs were grown on LB medium supplied with appropriate antibiotics for 24 h. Cells were harvested and resuspended to final OD_600_ of either 0.4 (all constructs other than RipH2) or 0.8 (RipH2) in infiltration medium (10 mM MgCl_2_, 10 mM MES pH 5.6). A blunt syringe was used to infiltrate the abaxial surface of 5-week old *N. benthamiana* or *N. tabacum* leaves. The symptoms of hypersensitive response were observed and photographed at 4 days post infiltration (dpi) for *N. benthamiana* or 6-7 dpi for *N. tabacum*.

Ion leakage assays were performed after infiltration of the same AGL1 strains into *N. benthamiana* or *N. tabacum* leaves. Plants were infiltrated with bacterial inoculum (OD_600_ 0.4) and two leaf discs of 9 mm diameter each were harvested daily from 0 to 6 dpi. The leaf discs were placed in 2 ml of distilled water (Millipore MilliQ system, Bedford MA) with shaking at 150 rpm for 2 h at room temperature. Ion leakage was measured using a conductivity meter (LAQUAtwin EC-33, Horiba) by loading 60 μl from each sample well. 3-4 technical replicates were used for each treatment. Statistical differences between treatments was assessed using Students two-tailed T-test with a *p*-value cutoff of 0.05.

## Results and Discussion

### Diversity and distribution of *R. solanacearum* phylotype I isolates across South Korea

Assemblies of thirty isolates of *R. solanacearum* collected from pepper and tomato in South Korea between 1999 and 2008 were generated from paired-end Illumina sequencing. Hybrid assemblies were generated with additional Nanopore sequence data for five isolates (Table 1). SPAdes Illumina assemblies were on average 5.7 Mb long, with scaffold counts ranging from 200 (Pe_26) to 306 (To_42). Unicycler hybrid assemblies for five isolates (Pe_1, Pe_3, Pe_27, Pe_39 and Pe_57) were on average 5.85 Mb in length across 2-9 scaffolds. Pe_39 and Pe_57 hybrid assemblies are considered complete as they are comprised of 2-4 circularized replicons.

Maximum likelihood trees were built using non-recombinant core genome alignments generated by readmapping to each Pe_57 replicon as reference. Recombination events identified by ClonalFrameML were removed to obtain non-recombinant trees of South Korean phylotype I strains (Fig. 1). Phylogenies produced using either the chromosome or megaplasmid as reference share nearly identical topologies, with four major clades of isolates and a single representative of a divergent lineage sampled in the eastern mainland (Pe_57). The chromosomal tree is slightly more resolved and has shorter branches leading to clades 1 and 2 than the megaplasmid tree. The two larger clades include strains isolated from both hosts: clade 2 includes isolates from central and northern regions while clade 4 includes spanning the entire country, including Jeju Island. A sister group to clade 2 includes solely pepper isolates (Pe_26, Pe_28, Pe_39 and Pe_1), while a smaller sister group to clade 4 includes a pair of tomato isolates (To_53, To_22). Other than these small clades, there is no strong phylogenetic signal of host of isolation: tomato and pepper isolates are distributed across the phylogeny. Although all but one tomato isolate was sampled from a single northern province and pepper isolates were sampled from seven other provinces, the tomato isolates show no evidence of phylogeographic clustering. Almost every province sampled has isolates from at least two clades, indicating diverse phylotype I isolates are circulating across many agricultural regions in South Korea.

**Figure 1.**
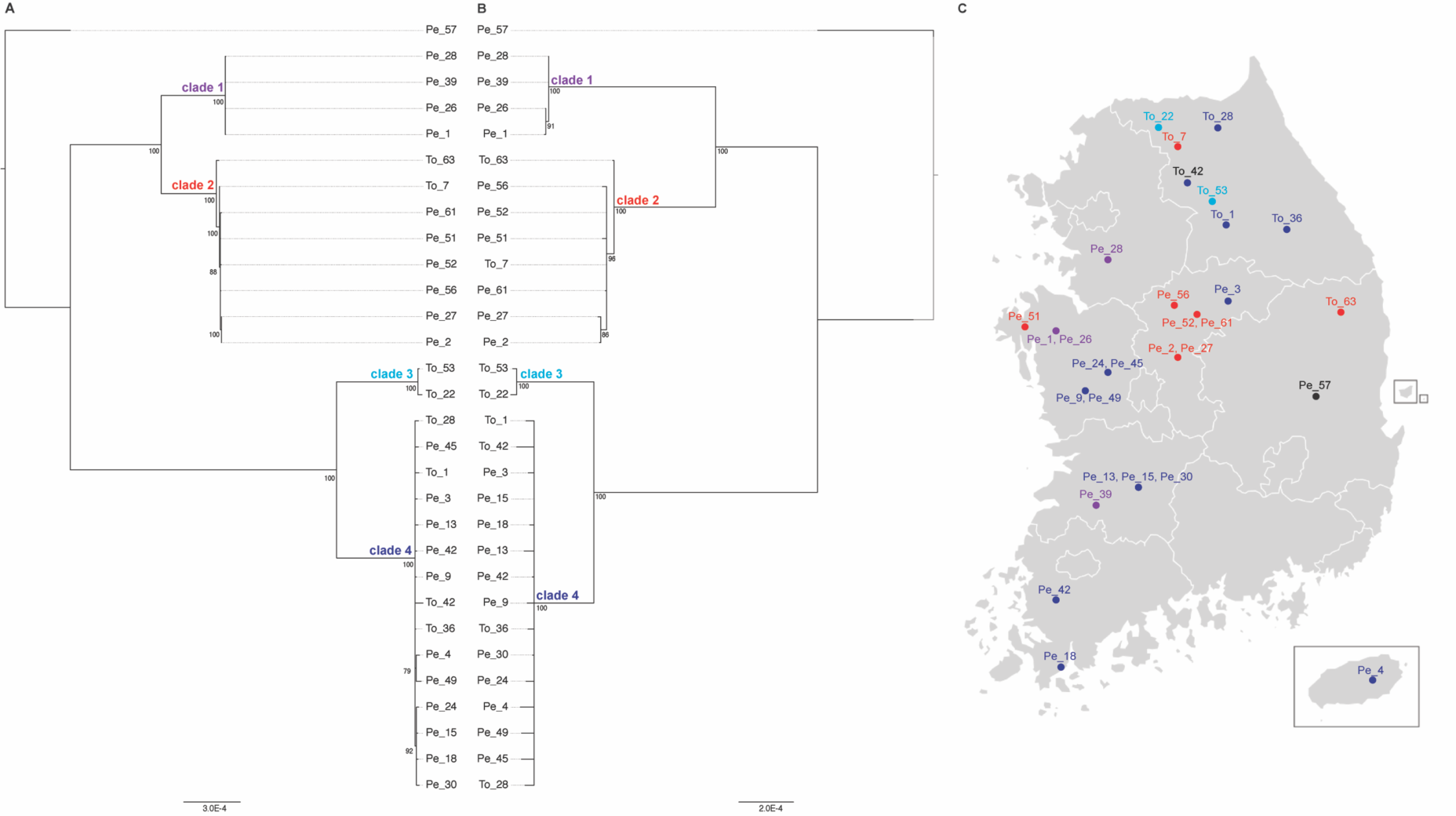
Diversity and distribution of South Korean phylotype I isolates. (A) Non-recombinant core chromosomal tree and (B) non-recombinant core megaplasmid tree of *R. solanacearum* phylotype I strains isolated from tomato (To) and pepper (Pe) hosts across South Korea (C). Core alignments were generated by readmapping to separate Pe_57 reference replicons. Phylogenetic trees were built with RAxML v7.2.8 and visualized using FigTree v1.4.3 displaying only the branches with bootstrap support score above 70. Node labels indicate the bootstrap support.

### Recombination of genes linked with virulence and competition

The cumulative impact of recombination on South Korean phylotype I isolates is enormous: 64.2% (2,026,835) of the gap-free non-mobile 3,155,880bp chromosome and 89.0% (1,511,546) of the gap-free non-mobile 1,698,146bp megaplasmid alignments are predicted to be affected by recombination in one or multiple strains at some stage in their evolutionary history (Table 2). Recombination occurs at only a slightly reduced frequency relative to mutation on both replicons but is far more likely to introduce substitutions, with ratios of the effect of recombination/mutation at 4.3 and 5.4 for the chromosome and megaplasmid, respectively. Fewer non-recombinant variant sites were retained for the megaplasmid (614) than the chromosomal (4,260) core alignment.

**Table 2.**
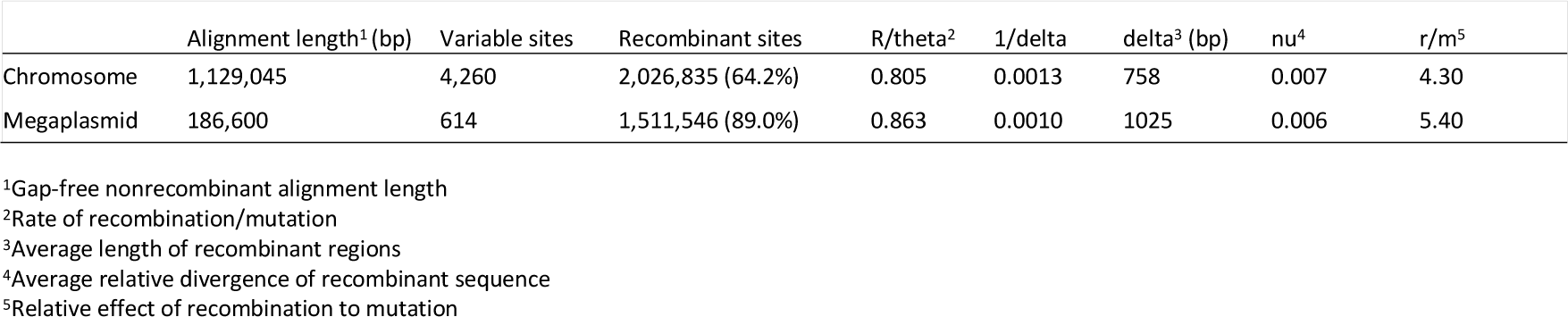
Impact of recombination on South Korean phylotype I isolates

The location of recombinant events within South Korean phylotype I genomes was assessed to determine whether genes affected by recombination are of relevance to pathogenesis in *R. solanacearum*. Extensive recombination events are present at ancestral nodes in the tree, while the range of predicted recent strain-specific recombination events varies between single recombination events for Pe_15 and Pe_28 to 27 recombination events across both replicons for To_42 and To_63. Since Pe_57 is a unique representative of a separate lineage in the phylogeny, many apparent strain-specific recombination events (118 and 77 on the chromosome and megaplasmid, respectively) are likely to have occurred earlier in the history of this lineage. There are distinct hotspots of recombination on both replicons (Fig. 2 and Table S2). 62 chromosomal and 46 megaplasmid genes are affected by two or more recent recombination events, some of which encompass multiple coding sequences (Table S2).

**Figure 2.**
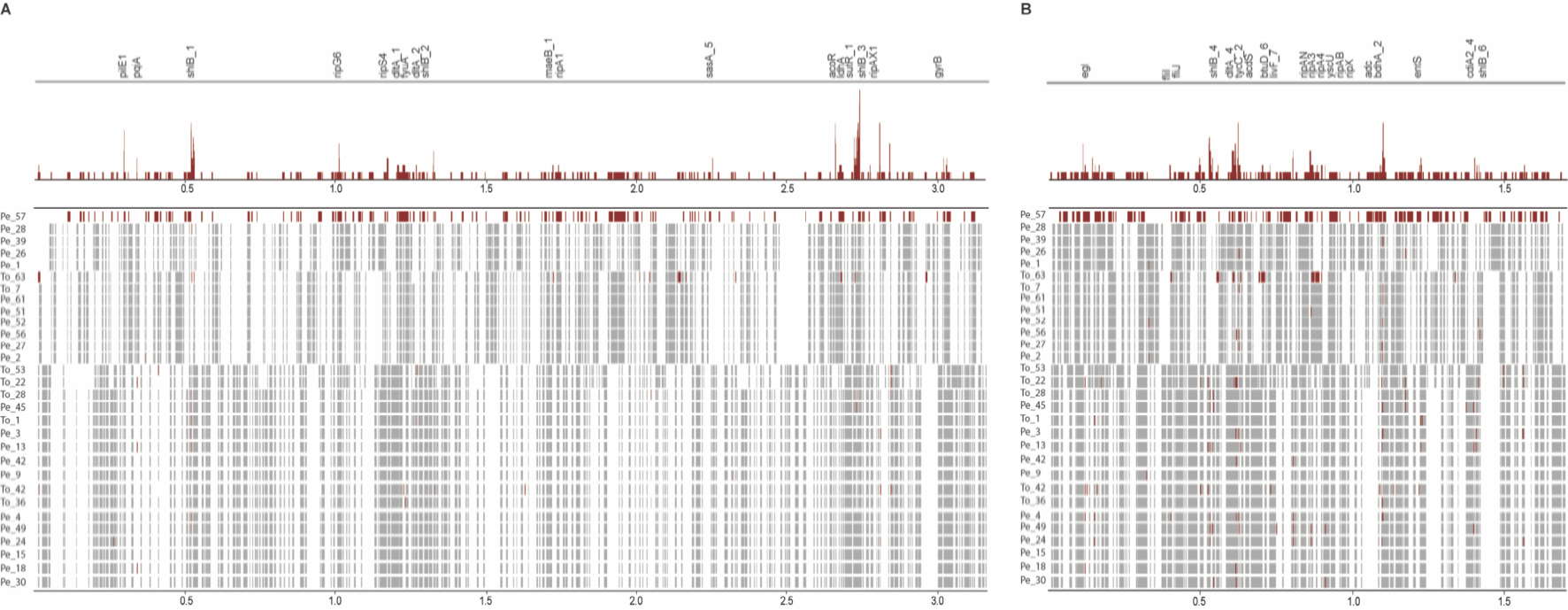
Recombination in South Korean phylotype I isolates. Recombination events identified by ClonalFrameML in chromosomal (A) and megaplasmid (B) sequences of South Korean *R. solanacearum* strains. Ancestral recombination events are shown in grey, and recent/strain-specific recombination events shown in red. Sum of strain-specific events shown at the top of each panel. Genes with two or more strain-specific events have gene IDs listed (hypothetical proteins not listed).

Many genes subject to recent recombination events are linked with virulence and microbial competition. Strikingly, six of these recombination hotspots overlap with six out of seven clusters of hemagglutinin and hemolysin (*shlB*) genes in Pe_57 (Fig. 2). Some of these clusters appear to be contact-dependent inhibition (CDI) loci, whose products mediate self/non-self-recognition and direct interference competition [42–44]. Contact-dependent inhibition is the product of two-partner (type V) secretion systems that assemble outer-membrane proteins (CdiB) involved in the export and presentation of large, toxic CdiA effectors on the cell surface [45]. CdiA effector protein binding to outer membrane receptors is followed by the transfer of the C-terminal toxin domain (CdiA-CT) into the susceptible bacterial cell cytoplasm, resulting in cell death. A cognate immunity protein (CdiI) confers resistance to toxin production by neighboring sister cells by binding and neutralizing specific CdiA-CT toxins; CdiI does not confer immunity to heterologous CdiA-CT toxins. CdiA-CT and CdiI sequences are highly variable both within and between species. Two clusters in Pe_57 (*shlB_1* and *shlB_4*) have *cdiA-cdiI-cdiO-cdiB*(*shlB*) arrangements typical of CDI loci in *Burkholderia*, as well as small ORFs and IS elements between *cdiI* and *cdiB*. Comparison of the *shlB_4* clusters in isolates for which nanopore assemblies are available (Pe_57, Pe_39, Pe_1, Pe_27 and Pe_3) reveals Pe_3 diverges significantly from other strains in the C-terminal domain of CdiA and cognate CdiI protein (Fig. S1). Recombination at CDI loci can be a mechanism promoting the maintenance of immunity between otherwise divergent isolates (e.g. Pe_57 vs Pe_39, Pe_1 and Pe_27) or a means of promoting antagonism via rapid diversification in toxin and cognate immunity proteins (e.g. Pe_27 vs Pe_3). The presence of multiple CDI loci in a single strain suggest both processes may operate simultaneously.

Recombination targets additional genes involved in microbial interactions and virulence. Up to eleven strain-specific recombination events were identified in genes annotated as D-alanine-polyphophoribitol ligase subunit 1 (*dltA_4*) and tyrocidine synthase *tycC_2* genes by Prokka. The closest BLASTn hit for this region in the phylotype I isolate GMI1000 is to a non-ribosomal peptide synthase locus called ralsomycin (*rmyA* and *rmyB*) or ralstonin which is linked with *R. solanacearum* invasion of fungal hyphae and the formation of chlamydospores [46, 47]. Five chromosomal (*ripA1, ripB, ripG6, ripG7, ripTAL*) and three megaplasmid (*ripH2, ripD, ripAW*) effectors show evidence of two or more strain-specific recombination events among the South Korean phylotype I isolates. Flagellar genes (*fliI* and *fliJ*), a type IV prepilin-like protein (*plpA*, annotated by Prokka as *pilE1*), a type III secretion system translocation protein (*yscU*) and component of the export pathway for enterobactin (*entS*), a high-affinity siderophore are also affected by multiple strain-specific recombination events in South Korean phylotype I isolates. The detection of recombination within gyrase B and endoglucanase is particularly notable given these are frequently used to infer relationships between *Ralstonia* strains.

### Core gene tree of the *R. solanacearum* species complex

In order to place the new South Korean strains and T3E dynamics in context of the structure of *R. solanacearum* species complex, a maximum-likelihood phylogeny was inferred from a concatenated alignment of core genes identified in an OrthoFinder pangenome analysis (Fig. 3). The core genome of the 30 South Korean phylotype I genomes and 68 NCBI genomes is represented by 1,680 single-copy orthogroups; 17.9% of the species complex pangenome. The *R. solanacearum* species complex has a large flexible genome, with 39.7% (3,720/9,366) of orthogroups present in 15 to 97 genomes and 40.1% of orthogroups in the cloud genome (1 to 15 genomes).

**Figure 3.**
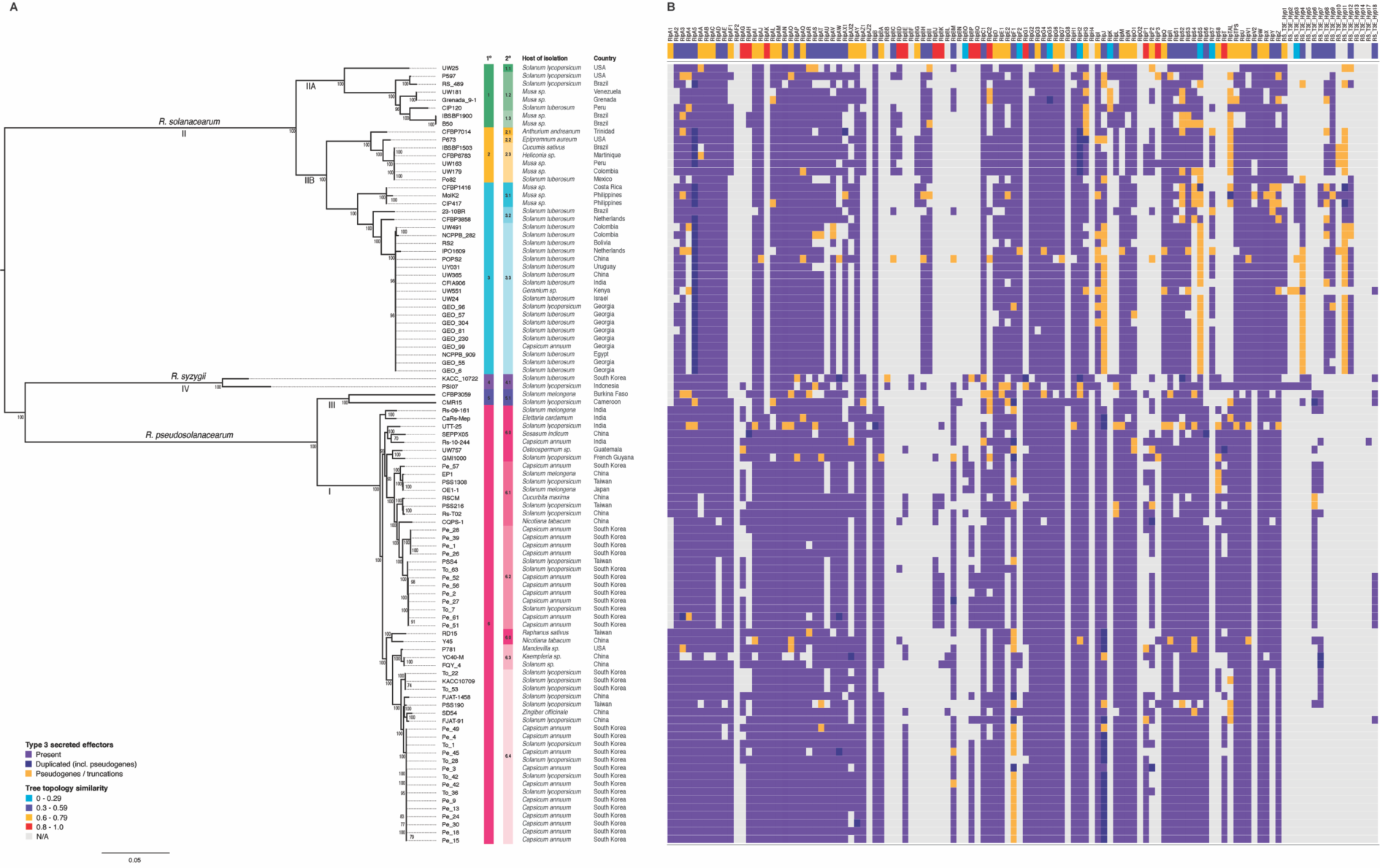
Phylogeny and type III secreted effector repertoires of the *R. solanacearum* species complex. (A) Maximum likelihood tree inferred using a non-recombinant core gene alignment of single-copy orthologs. Species and phylotype designation shown above and below branches, respectively. Nodes with bootstrap support values under 70 collapsed. Primary (1°) and secondary (2°) cluster assignment by hierBAPS shown next to strain labels. (B) Type III effector repertoires. Top row displays normalized Robinson Fould’s distance between the effector family gene tree and core gene tree shown at left. Lower values (e.g. blue) indicate greater topological similarity between effector and core gene tree.

The four major phylotypes (and three separate species) are displayed in the core gene tree. The first level of population clustering is consistent with phylotype designations I, IIA, III and IV, while phylotype IIB falls into two separate clusters. Clonal expansion of multiple groups is evident: a phylotype IIB subclade (3.3) typically associated with *S. tuberosum* infections has been sampled from three continents (South America, Africa and Eurasia). The two larger clades of South Korean phylotype I isolates correspond to subclades 6.2 and 6.4, primarily sampled from solanaceous hosts in China, Taiwan and South Korea (Fig. 3). The divergent pepper isolate Pe_57 falls into a cluster of East Asian isolates from different hosts: eggplant (EP1, OE1-1); tomato (PSS1308, PSS216, Rs-To2); tobacco (CQPS-1) and squash (RSCM).

### Distribution of type III secreted effectors across the RSSC

The distribution of type III secreted effectors across the RSSC was determined to gain an understanding of the extent of variation within Korean isolates and place these in the context of the variation in the RSSC as a whole (Fig. 3B). The average T3E complement varies between phylotypes: phylotype I has on average 71 effector genes per strain, while phylotype II has an average of 61 effector genes per strain. Phylotypes III and IV are each represented by a pair of strains: these have an average of 58 and 64 T3Es per strain respectively. Many T3E families are broadly distributed across the entire species complex: 64 out of 118 (54%) of T3Es are present in at least half of the isolates included here. *RipAB* is present in all strains but YC40-M (*Kaempferia* sp., China) and *ripU* is potentially pseudogenised in CFBP3059 (*Solanum melongena*, Burkina Faso) but is otherwise ubiquitous. There are some highly amplified T3E families present across much of the species complex, like *ripA* and *ripB*. Diversification of these families may be linked with host recognition; the broad distribution suggests they encode critical virulence or host subversion functions.

There are both clade- and phylotype-specific patterns of T3E presence and absence. A set of fourteen T3Es (*ripA1, ripAG, ripAK, ripBA, ripBE, ripBJ, ripBK, ripBO, ripBP, ripG1, ripS6, ripS8, ripT* and *Hyp6*) are present in one or more phylotype I strain, but are absent from phylotype II. This is likewise the case for *ripAH*, though it is present in two phylotype II strains. Three T3Es (*ripJ, ripS5* and *ripTAL*) are mostly truncated or pseudogenized in phylotype II, while full-length versions are observed in phylotype I. Conversely, seven T3Es (*ripBC, ripBG, ripBH, ripBI, ripK, ripS7* and *ripV2*) and many hypothetical T3Es (*Hyp1, Hyp3, Hyp4, Hyp8, Hyp9, Hyp11* and *Hyp12*) are present in phylotype II but absent in phylotype I.

There are some apparent patterns of recurrent T3E loss across phylotypes I and II. In phylotype II these losses are neither clade nor host-specific, affecting IPO1609 (*S. tuberosum*, Netherlands), Po82 (*S. tuberosum*, Mexico), RS2 (*S. tuberosum*, Bolivia), UW551 (*Geranium sp*., Kenya), UW25 (*S. lycopersicum*, USA), POPS2 (*S. lycopersicum*, China), RS489 (*S. lycopersicum*, Brazil). The apparent losses in phylotype I impact tomato strains PSS190 from Taiwan, UTT-25 from India, KACC10709, To_22 and To_53 from South Korea. In addition, Cars-Mep strain isolated from *Elettaria cardamom* from India shows significant effector losses. These patterns should be interpreted with caution as some apparent losses (or indeed truncations and pseudogenizations) may be due to variable sequencing and assembly quality.

Despite their ubiquity across the species complex, T3Es are nevertheless subject to recombination. Individual T3E gene trees were compared with the core gene tree to determine whether they exhibit evidence of horizontal transfer. The normalized Robinson Fould’s (nRF) distance implemented in ETE3 was used as a measure of topological similarity between individual T3E trees and the core gene tree. Some T3Es, including *ripAK, ripBD, ripAH, ripP1, ripC2, ripAG, ripG1* and *ripT* have divergent tree topologies (nRF > 0.8), indicating they have been subject to dynamic exchange throughout their evolutionary histories. T3Es with topologies consistent with the core gene tree are typically limited to specific clades (e.g. *ripK, ripBO, ripF2* and *ripS7*, with nRF < 0.2). There are however three exceptions: *ripG5, ripH2* and *ripS5* are broadly distributed across the RSSC and are less frequently exchanged, though *ripS5* is pseudogenized in many phylotype II strains. Allelic variation is likely to determine fine-scale differences in host range among T3Es with broad distributions across the *Ralstonia* species complex.

### Allelic diversity in T3Es impacts host recognition

Allelic diversification of T3Es is likely to play an important role in host adaptation. The link between defense induction and allelic diversity among South Korean isolates was examined for six T3Es: *ripA1, ripAA, ripAY, ripE1, ripA5* and *ripH2*. These families were chosen as they were reported to trigger programmed cell death when overexpressed in *Nicotiana* spp. [18–22]. Alleles were classified into groups on the basis of shared amino acid sequence identity; patterns of diversity broadly reflected the phylogenetic groupings in the non-recombinant core genome tree for South Korean strains phylotype I strains (Fig. 4A, B).

**Figure 4.**
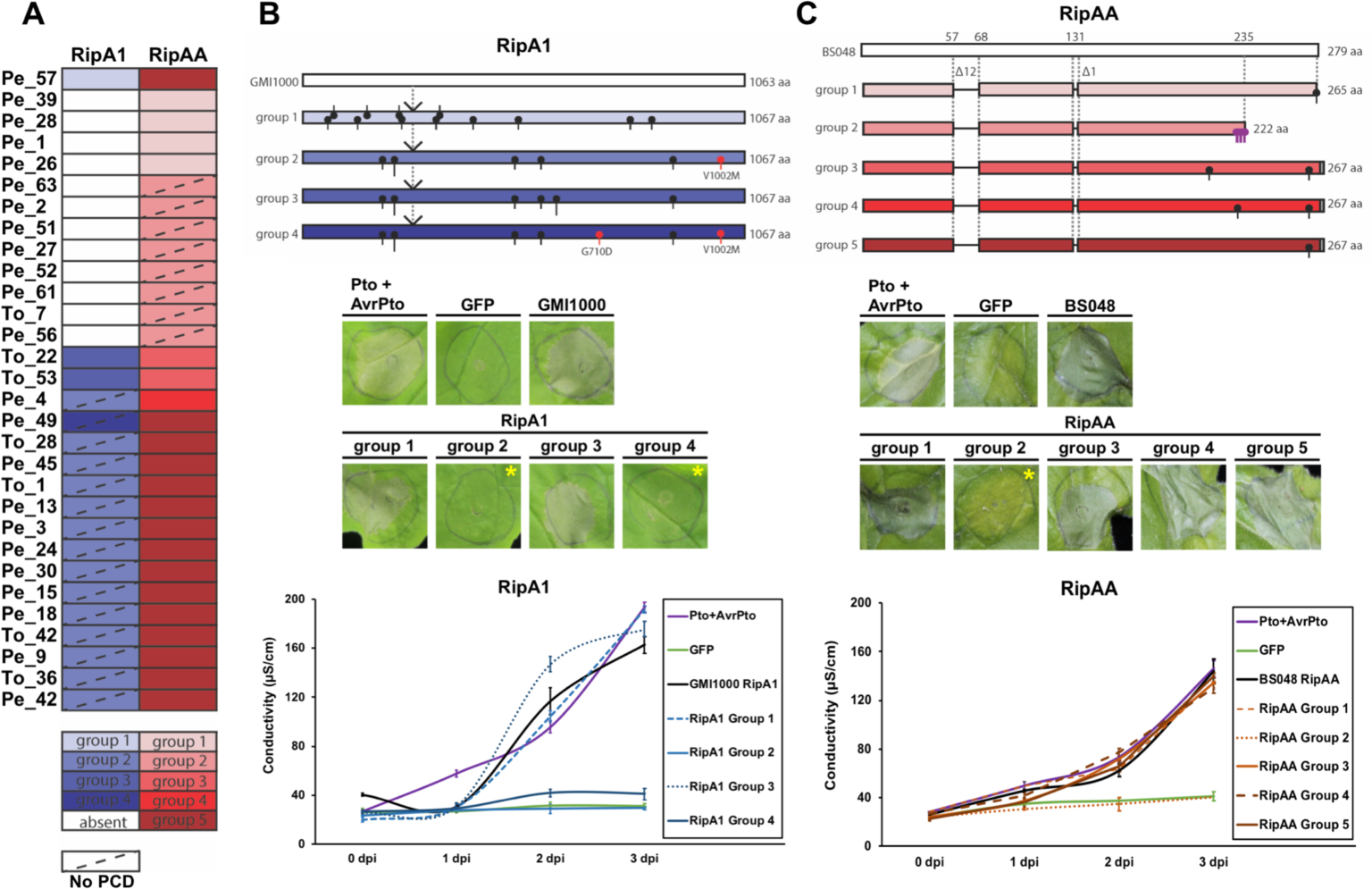
Effector alleles of RipA1 and RipAA display variable recognition responses in *N. benthamiana*. (A) Effector alleles were classified into groups (A) based on amino acid sequence alignments (B) and representatives of each group were tested for their ability to trigger programmed cell death (PCD, shown in C) and increased ion leakage (D), markers of host recognition. *Agrobacterium* strains expressing allelic variants were infiltrated into *N. benthamiana* leaves at OD_600_ 0.4 and PCD was photographed 3 days post infection (dpi). Asterisks in (C) indicate absence of PCD for RipA1 allele groups 2 and 4, and RipAA group 2. GFP is a negative control, Pto+AvrPto, GMI1000 RipA1 and BS4048 RipAA refer to positive controls. Ion leakage in (D) was measured at 0, 1, 2 and 3 dpi and plotted as average conductivity of 4 technical replicates, error bars represent standard error of mean (SEM).

Representative alleles from each group (Table S3) were transiently expressed in *N. benthamiana* and *N. tabacum* plants by agroinfiltration in order to score their ability to elicit a programmed cell death and ion leakage, markers of host recognition. Overexpression of RipA1 group 2 and group 4 alleles failed to elicit cell death, while group 1 and group 3 alleles triggered cell death and ion leakage (Fig. 4C, D). One of two substitutions present solely in groups 2 and 4 is likely responsible for the loss of recognition (V1002M); an additional G710D substitution is present in group 4. Host recognition of RipAA alleles varied as well: a single allele (group 2) failed to elicit cell death. A C-terminal truncation preceded by multiple substitutions is likely responsible for the loss of recognition of group 2 RipAA in *N. benthamiana* (Fig. 4C, D). We found RipA1 and RipAA protein variants accumulated to significant levels *in planta*, suggesting the phenotypes observed are not due to variation in protein stability but rather the loss of host recognition (Fig. S2). Both RipAY alleles present among South Korean phylotype I strains trigger weak cell death and intermediate ion leakage, while all RipE1 and RipA5 alleles caused strong cell death and ion leakage (Fig. S3). In contrast, no RipH2 alleles triggered cell death and ion leakage when overexpressed in *N. tabacum* (Fig. S3).

## Conclusions

*Ralstonia* infections of staple carbohydrate and vegetable crops threaten regional food security and agricultural productivity. The 30 South Korean isolates shown here encompass much of the diversity of phylotype I. Phylotype I is likely to have been present in East Asia for some time, though recent clonal expansions and dissemination between agricultural regions is apparent among South Korean isolates. High rates of recombination among the *R. solanacearum* species complex enhance the potential for rapid adaptation and the emergence of novel pathogen variants. Recombination affecting multiple virulence-related genes was identified among the South Korean strains. Hotspots of recombination overlap with contact-dependent inhibition loci, indicating microbial competition plays a critical role in *Ralstonia* evolution, perhaps during stages in its lifecycle when it must compete for nutrients in soil and freshwater environments.

The *R. solanacearum* species complex has an expanded T3E repertoire relative to other bacterial plant pathogens. Many T3Es have evolutionary histories inconsistent with the core gene topology, revealing how recombination can reshuffle virulence factor repertoires. Though some effector families may be less frequently horizontally transferred than others, host adaptation and selection imposed by host recognition can result in allelic variation linked with altered host specificity. The integration of pathogen population genomics with molecular plant pathology allows for the identification of conserved pathogen virulence proteins with consistent recognition and elicitation activity, providing suitable targets for breeding crops with durable resistance to *R. solanacearum* phylotype I in South Korea.

## Acknowledgements

We gratefully acknowledge support for this work from the following sources: Next-Generation BioGreen 21 Program (Plant Molecular Breeding Center No. PJ01317501), Rural Development Administration (RDA) and the National Research Foundation (NRF) of Korea grants funded by the Korea government (MSIT) (NRF-2019R1A2C2084705 & 2018R1A5A1023599 (SRC)), Republic of Korea (K.H.S.); Creative-Pioneering Researchers Program through Seoul National University (C.S.); Royal Society of New Zealand Marsden Fast-Start (MAU1709) and support from the Max Planck Society (H.C.M.). We thank Sven Kuenzel (MPI for Evolutionary Biology) for Illumina sequencing.

## Conflicts of Interest

The authors declare that there are no conflicts of interest.

## Figures

**Figure S1.**
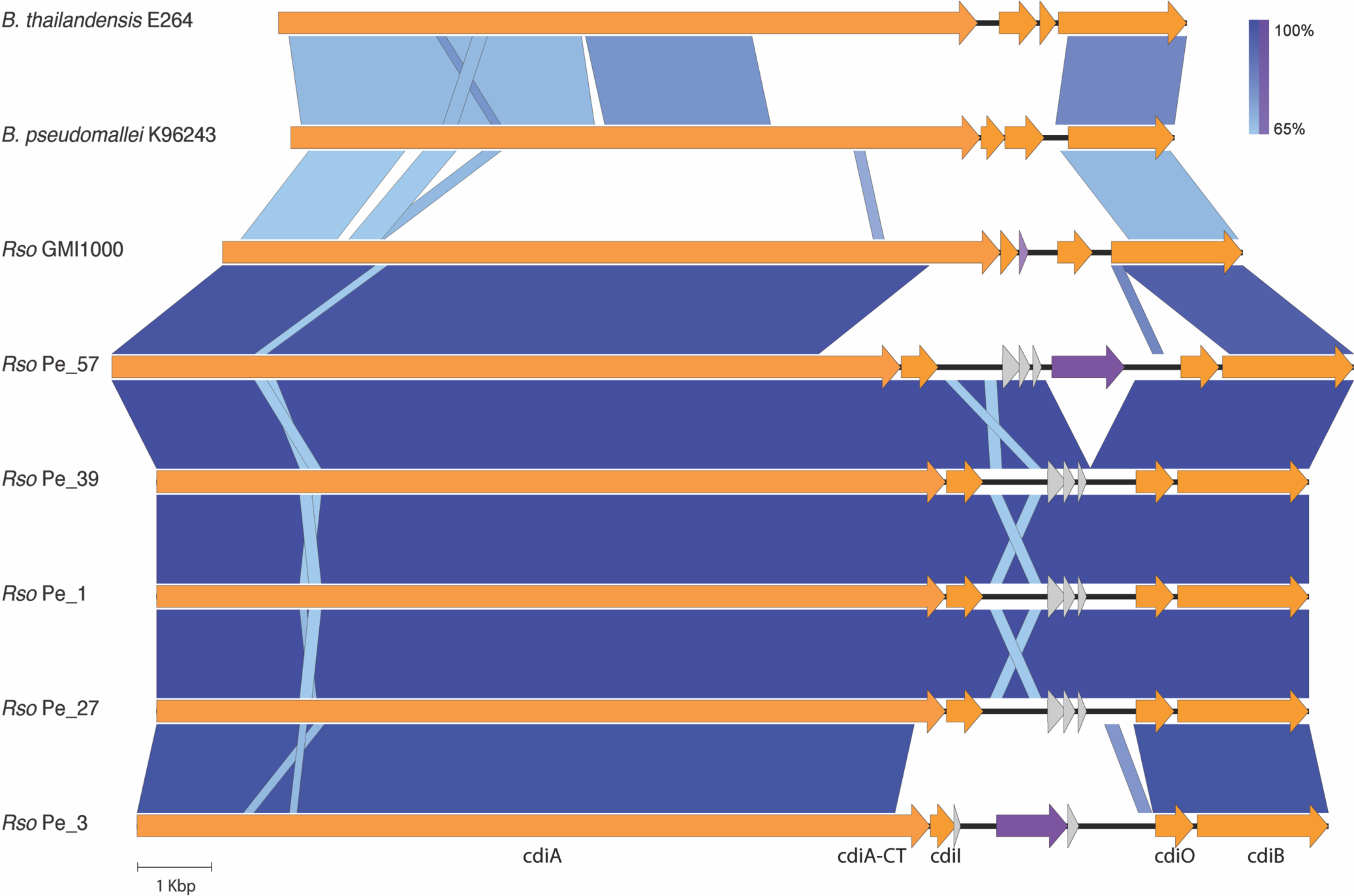
Contact-dependent inhibition loci in *Ralstonia* phylotype I isolates. CDI regions homologous to the shlB_4 region in Pe_57 were identified in nanopore assemblies of South Korean phylotype I isolates as well as the complete genome of GMI1000, *B. thailandensis* E264 and *B. pseudomallei* K96243. Orange CDS refer to (left to right) *cdiA, cdiI, cdiO* and *cdiB*; grey CDS refer to hypotheticals and purple CDS refer to IS5-family transposases IS1021 (Pe_57) and ISRso18 (Pe_3). The small light purple CDS is a pseudogenized IS (GMI1000). The cdiA-CT domain is immediately adjacent to cdiI. Vertical blocks indicate regions of shared similarity shaded according to BLASTn similarity.

**Figure S2.**
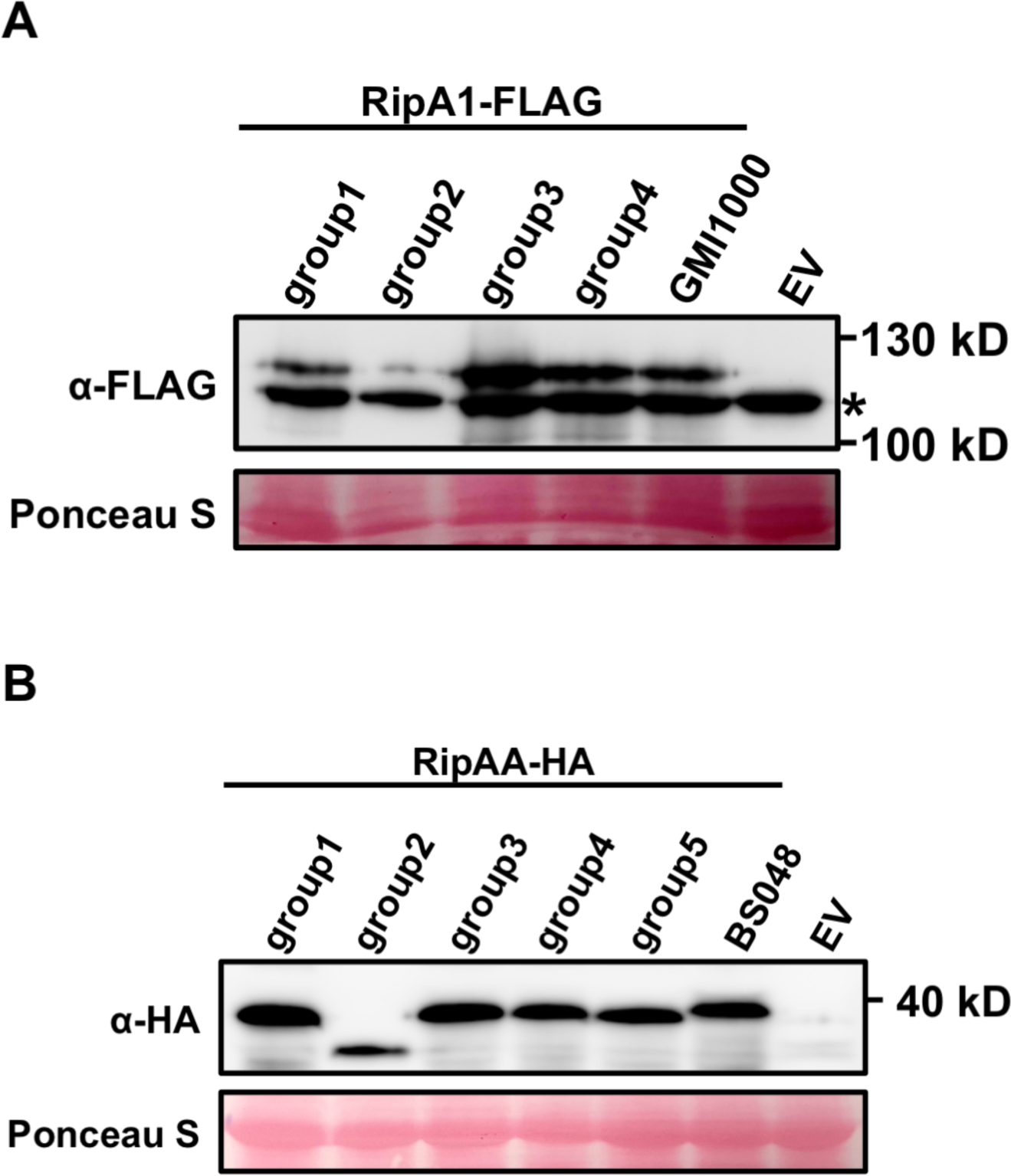
Loss of host recognition is not a product of differential protein accumulation *in planta*. RipA1 (A) and RipAA (B) protein accumulation does not explain the loss of programmed cell death phenotype when overexpressed in *N. benthamiana* leaves. Proteins were overexpressed in *N. benthamiana* leaves using agroinfiltration (OD_600_ 0.4). Proteins were sampled at 2 day post infiltration and subjected to western blot analysis to visualize their accumulation using anti FLAG and anti HA antibodies. A nonspecific band present in the RipA1 immunoblot is denoted with asterisk. Ponceau S staining provided to confirm equal amounts of protein loaded.

**Figure S3.**
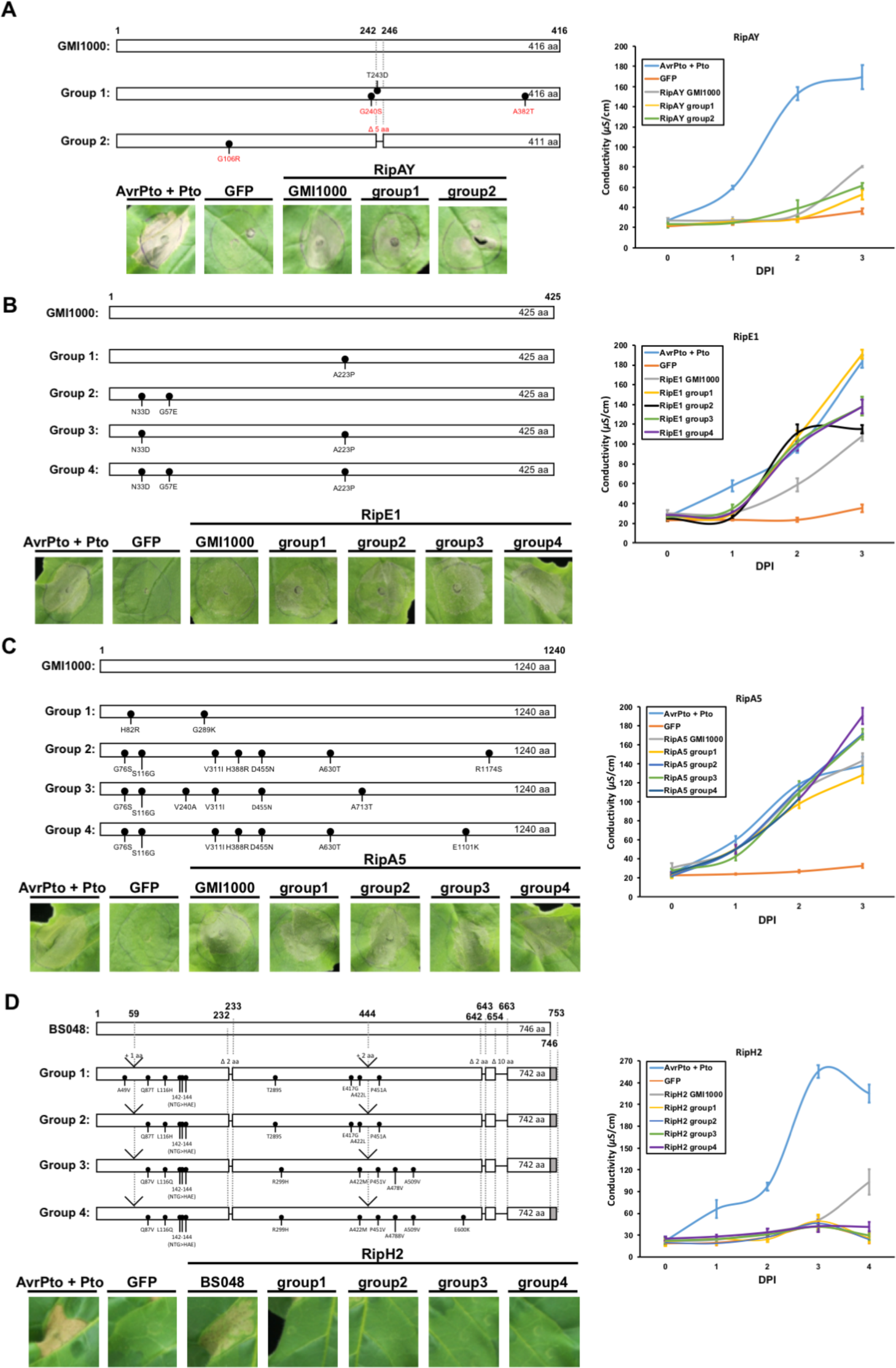
Consistent recognition of T3E alleles in *R. solanacearum* phylotype I isolates. RipAY (A), RipE1 (B), RipA5 (C) and RipH2 (D) were overexpressed in *N. benthamiana* (A,B,C) or *N. tabacum* (D) leaves and PCD symptoms photographed at 4 dpi. Ion leakage was quantified from the infiltrated tissue. Average of 4 technical replicates are plotted on the graph with error bars representing S.E.M.

**Table S1.** Publicly available *RSSC* assemblies included in this work.

| Strain | Phylotype | Host of isolation | Location of isolation | WGS Project | Genbank Accession |
| --- | --- | --- | --- | --- | --- |
| GMI1000 | I | *Solanum lycopersicum* | Kourou, French Guyana |  | GCA_000009125.1 |
| Rs-09-161 | I | *Solanum melongena* | Goa, India | JHBO01 | GCA_000671335.1 |
| CaRs-Mep | I | *Elettaria cardamum* | Kerala, India | MCBM01 | GCA_001855495.1 |
| UW757 | I | *Osteospermum* | Escuintla, Guatemala | LFJP01 | GCA_001645725.1 |
| UTT-25 | I | *Solanum lycopersicum* | India | PPFC02 | GCA_002930085.2 |
| Rs-10-244 | I | *Capsicum annuum* | Andaman Islands, India | JHAM01 | GCA_000671315.1 |
| SEPPX05 | I | *Sesamum indicum* | China |  | GCA_002162015.1 |
| EP1 | I | *Solanum melongena* | Guangdong, China |  | GCA_001891105.1 |
| PSS1308 | I | *Solanum lycopersicum* | Taiwan | MOLO01 | GCA_001870805.1 |
| OE1-1 | I | *Solanum melongena* | Japan |  | GCA_001879565.1 |
| RSCM | I | *Cucurbita maxima* | Guangdong, China |  | GCA_002894285.1 |
| PSS216 | I | *Solanum lycopersicum* | Hsinchu, Taiwan | MOLJ01 | GCA_001876975.1 |
| Rs-T02 | I | *Solanum lycopersicum* | Guangxi, China | LKVH01 | GCA_001484095.1 |
| CQPS-1 | I | *Nicotiana tabacum* | Chongqing, China |  | GCA_002220465.1 |
| PSS4 | I | *Solanum lycopersicum* | Taiwan | MOLK01 | GCA_001876985.1 |
| RD15 | I | *Raphanus sativus* | Taiwan | MNCM01 | GCA_001854265.1 |
| Y45 | I | *Nicotiana tabacum* | Yunnan, China | AFWL01 | GCA_000223115.2 |
| P781 | I | *Mandevilla sp.* | Florida, USA | JXLK01 | GCA_001644865.1 |
| FQY_4 | I | *Solanum sp.* | Guizhou, China |  | GCA_000348545.1 |
| YC40-M | I | *Kaempferia sp.* | Shajiang, China |  | GCA_001663415.1 |
| KACC10709 | I | *Solanum lycopersicum* | South Korea |  | GCA_001708525.1 |
| SD54 | I | *Zingiber officinale* | China | ASQR02 | GCA_000430925.2 |
| FJAT-91 | I | *Solanum lycopersicum* | Fujian, China |  | GCA_002155245.1 |
| FJAT-1458 | I | *Solanum lycopersicum* | China |  | GCA_001887535.1 |
| PSS190 | I | *Solanum lycopersicum* | Taiwan | MOLP01 | GCA_001870825.1 |
| UW25 | IIA | *Solanum lycopersicum* | USA | NCTK01 | GCA_002251695.1 |
| CIP120 | IIA | *Solanum tuberosum* | Peru | JXAY01 | GCA_001644795.1 |
| RS_489 | IIA | *Solanum lycopersicum* | Brazil |  | GCA_002549815.1 |
| P597 | IIA | *Solanum lycopersicum* | USA | JIBY01 | GCA_001644805.1 |
| IBSBF1900 | IIA | *Musa sp.* | Brazil | CDRW01 | GCA_001373275.1 |
| B50 | IIA | *Musa sp.* | Brazil | CDMA01 | GCA_000825785.2 |
| Grenada_9-1 | IIA | *Musa sp.* | Grenada | CDLW01 | GCA_000825845.2 |
| UW181 | IIA | *Musa sp.* | Venezuela | CDSB01 | GCA_001373315.1 |
| CFBP7014 | IIB | *Anthurium andreanum* | Trinidad | CDRJ01 | GCA_001373255.1 |
| P673 | IIB | *Epipremnum aureum* | USA | JALO01 | GCA_000525615.1 |
| CFBP6783 | IIB | *Heliconia sp.* | Martinique | JXAZ01 | GCA_001644815.1 |
| IBSBF1503 | IIB | *Cucumis sativus* | Brazil |  | GCA_001587155.1 |
| Po82 | IIB | *Solanum tuberosum* | Mexico |  | GCA_000215325.1 |

**Table S1. Publicly available RSSC assemblies included in this work - *continued***
| Strain | Phylotype | Host of isolation | Location of isolation | WGS Project | Genbank Accession |
| --- | --- | --- | --- | --- | --- |
| UW179 | IIB | *Musa* sp. | Colombia | CDLZ01 | GCA_000825805.2 |
| UW163_2 | IIB | *Musa* sp. | Peru |  | GCA_001587135.1 |
| CFBP1416 | IIB | *Musa* sp. | Costa Rica | CDLX01 | GCA_000825925.2 |
| MolK2 | IIB | *Musa* sp. | Philippines | CAHW02 | GCA_000212635.2 |
| CIP417 | IIB | *Musa* sp. | Philippines | CDMD01 | GCA_000825825.2 |
| 23-10BR | IIB | *Solanum tuberosum* | Brazil | JQOI01 | GCA_000749995.1 |
| CFBP3858 | IIB | *Solanum tuberosum* | Netherlands | CDSB01 | GCA_001373315.1 |
| UW491 | IIB | *Solanum tuberosum* | Colombia | LVKU01 | GCA_001696845.1 |
| NCPPB_282 | IIB | *Solanum tuberosum* | Colombia | JQSH01 | GCA_000750575.1 |
| RS2 | IIB | *Solanum tuberosum* | Bolivia | CDRX01 | GCA_001373295.1 |
| IPO1609 | IIB | *Solanum tuberosum* | Netherlands | CDGL01 | GCA_001050995.1 |
| UW551 | IIB | *Geranium* sp. | Kenya | NCTI01 | GCA_002251655.1 |
| GEO_99 | IIB | *Capsicum annuum* | Georgia | MZNB01 | GCA_002029865.1 |
| UW24 | IIB | *Solanum tuberosum* | Israel | LVKT01 | GCA_001696855.1 |
| GEO_230 | IIB | *Solanum tuberosum* | Georgia | PHHL01 | GCA_002894795.1 |
| POPS2 | IIB | *Solanum tuberosum* | China | JQSI01 | GCA_000750585.1 |
| GEO_304 | IIB | *Solanum tuberosum* | Georgia | PHHO01 | GCA_002894775.1 |
| CFIA906 | IIB | *Solanum tuberosum* | India | MZNA01 | GCA_002029895.1 |
| GEO_96 | IIB | *Solanum lycopersicum* | Georgia | MZNA01 | GCA_002029895.1 |
| UW365 | IIB | *Solanum tuberosum* | China | LVKS01 | GCA_001696865.1 |
| GEO_6 | IIB | *Solanum tuberosum* | Georgia | PHHM01 | GCA_002894765.1 |
| NCPPB_909 | IIB | *Solanum tuberosum* | Egypt | JNGD01 | GCA_000710695.1 |
| GEO_57 | IIB | *Solanum tuberosum* | Georgia | MXAM01 | GCA_002029885.1 |
| GEO_55 | IIB | *Solanum tuberosum* | Georgia | PHHP01 | GCA_002894845.1 |
| UY031 | IIB | *Solanum tuberosum* | Uruguay |  | GCA_001299555.1 |
| GEO_81 | IIB | *Solanum tuberosum* | Georgia | PHHN01 | GCA_002894785.1 |
| KACC_10722 | IV | *Solanum tuberosum* | South Korea |  | GCA_001586135.1 |
| PSI07 | IV | *Solanum lycopersicum* | Indonesia |  | GCA_000283475.1 |
| CMR15 | III | *Solanum lycopersicum* | Cameroon |  | GCA_000427195.1 |
| CFBP3059 | III | *Solanum melongena* | Burkina Faso | JXBA01 | GCA_001644855.1 |

**Table S2.**
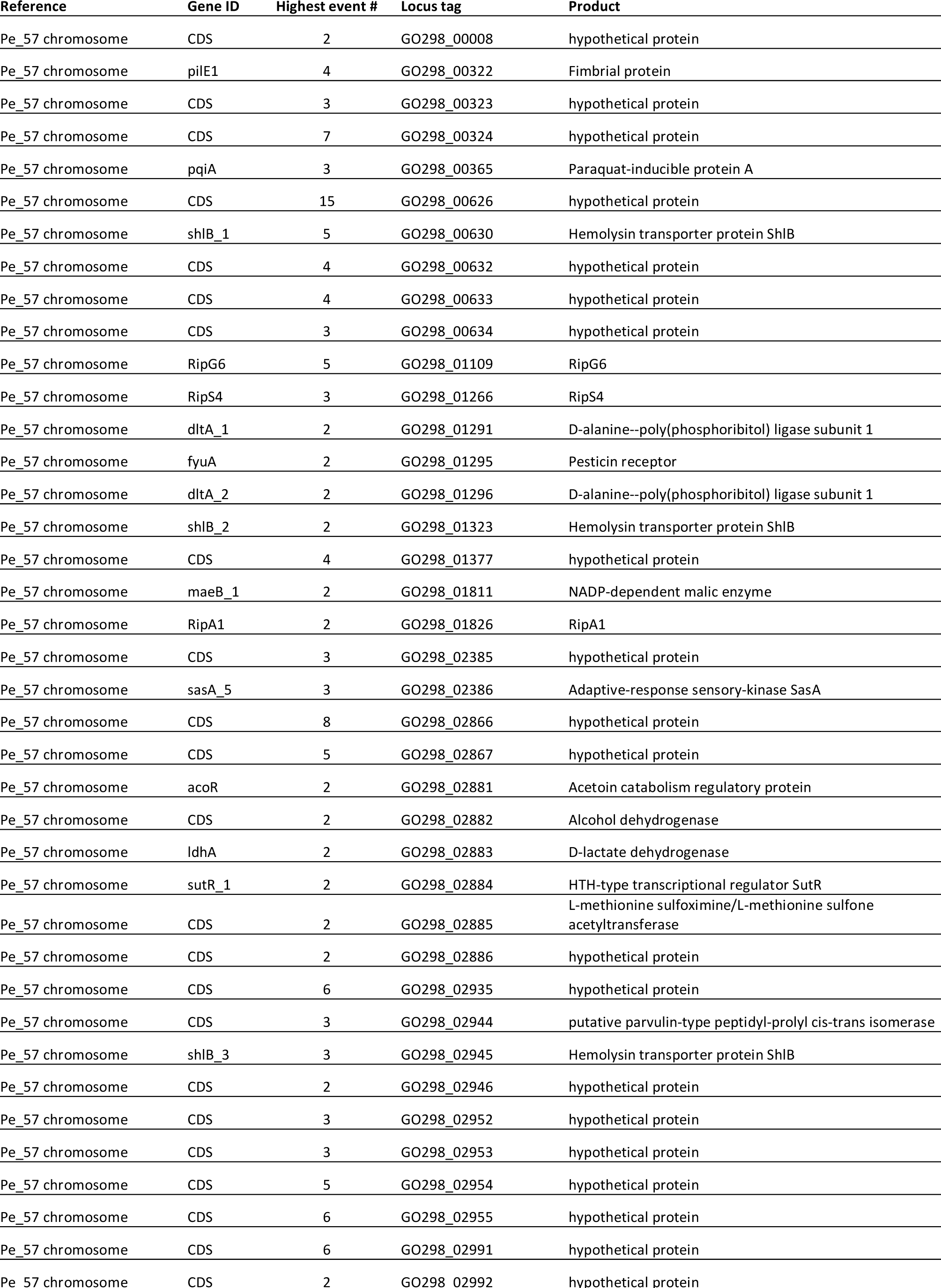
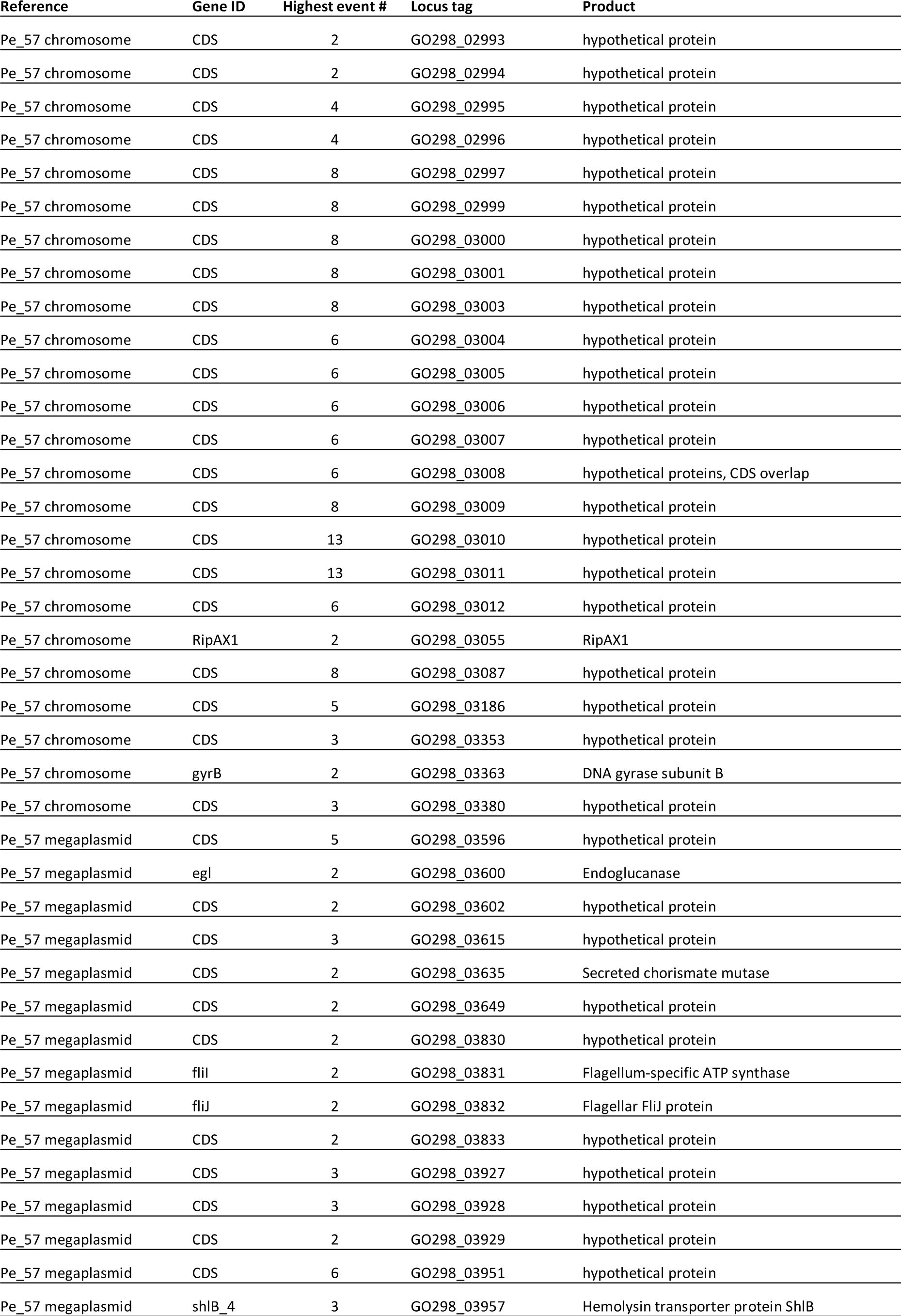
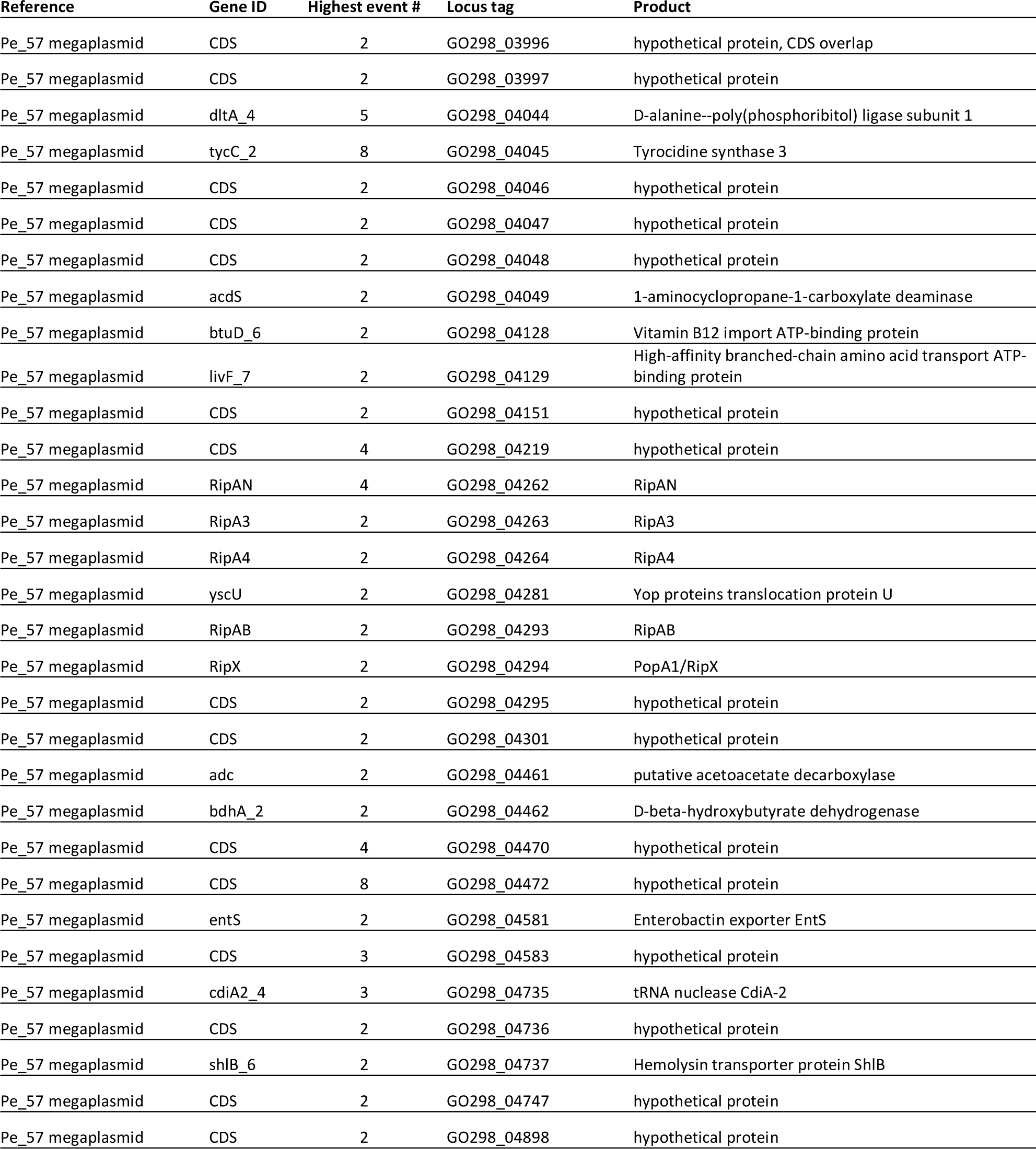
Genes with 2+ strain-specific recombination events.

**Table S3.**
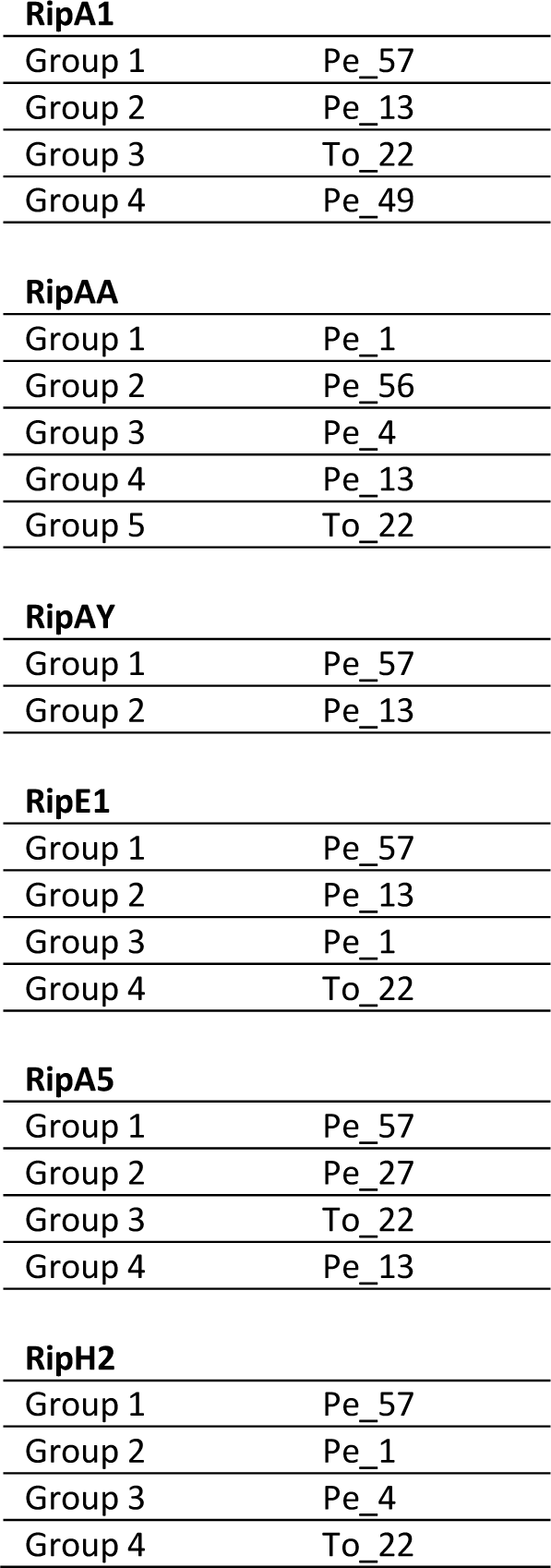
Strains used as a source of representative effector alleles for testing host recognition.

**RipA1**
|  |  |
| --- | --- |
| Group 1 | Pe_57 |
| Group 2 | Pe_13 |
| Group 3 | To_22 |
| Group 4 | Pe_49 |
| **RipAA** |  |
| Group 1 | Pe_1 |
| Group 2 | Pe_56 |
| Group 3 | Pe_4 |
| Group 4 | Pe_13 |
| Group 5 | To_22 |
| **RipAY** |  |
| Group 1 | Pe_57 |
| Group 2 | Pe_13 |
| **RipE1** |  |
| Group 1 | Pe_57 |
| Group 2 | Pe_13 |
| Group 3 | Pe_1 |
| Group 4 | To_22 |
| **RipA5** |  |
| Group 1 | Pe_57 |
| Group 2 | Pe_27 |
| Group 3 | To_22 |
| Group 4 | Pe_13 |
| **RipH2** |  |
| Group 1 | Pe_57 |
| Group 2 | Pe_1 |
| Group 3 | Pe_4 |
| Group 4 | To_22 |

## Notes

### Competing Interest Statement

The authors have declared no competing interest.

